# Elucidation of Genome Polymorphisms in Emerging SARS-CoV-2

**DOI:** 10.1101/2020.07.22.215731

**Authors:** Manisha Ray, Saurav Sarkar, Surya Narayan Rath, Mukund Namdev Sable

## Abstract

The COVID-19 pandemic is having a devastating effect on the healthcare system and the economy of the world. The unavailability of a specific treatment regime and a candidate vaccine yet opens up scope for new approaches and discoveries of drugs for mitigation of the sufferings of humankind due to the disease. The present isolated whole-genome sequences of SARS-CoV-2 from 11 different nations subjected to evolutionary study and genome-wide association study through *in silico* approaches including multiple sequence alignment, phylogenetic study through MEGA7 and have been analyzed through DNAsp respectively. These investigations recognized the nucleotide varieties and single nucleotide mutations/polymorphisms on the genomic regions as well as protein-coding regions. The resulted mutations have diversified the genomic contents of SARS-CoV-2 according to the altered nucleotides found in 11 genome sequences. India and Nepal have found to have progressively more distinct species of SARS-CoV-2 with variations in Spike protein and Nucleocapsid protein-coding sites. These genomic variations might be the explanation behind the less case fatality rate of India and Nepal dependent on the populaces. The anticipated idea of this investigation upgrades the information about genomic medication and might be useful in the planning of antibodies against SARS-CoV-2.

## 1. Introduction

The coronavirus disease 2019 (COVID-19), is caused by severe acute respiratory syndrome coronavirus-2 (SARS-CoV-2). It is the 7^th^ beta coronavirus isolated from Wuhan, China in December 2019^1^. COVID-19 is not only just threatening the public health but also affecting social stability and economic development^2^. Among the previously identified six human CoVs including 229E, HKU1, NL63, OC43, SARS, and MERS the highly pathogenic strains were SARS, MERS in contrast to the others^3^. Of them, COVID19 has pandemic since it has spread more than 200 countries rapidly^1^. The World Health Organization declared it as a public health emergency of international concern because of its rapid spread to so many countries^4^. Along with the respiratory illness, the strong virulence of SARS-CoV-2 also causes multi-organ failure, including renal and neurological ailments^2,5^.

CoVs are single-stranded RNA viruses, and share a large genome around 30kb, express abundant replica genes encoding non-structural proteins involving approximately 20kb of the genome^6^. The genome comprises of a 5’ terminal an open reading box (ORF) 1a/b coding region, Spike (S) glycoprotein, Envelope (E) protein, Membrane (M) protein, Nucleocapsid (N) protein and a 3’ terminal noncoding region. Among these S protein specifically bind to the receptor of the host cell and mediates membrane fusion and virus entry^5, 7^. In contrast, N protein has involved in the assembly of the virus inside the host cell, and it is necessary for viral RNA transcription and replication^7-8^; thus these two proteins have been reported as the most prominent proteins for progression of viral infections. According to the updated data of the WHO, the ratio of confirmed cases and the death rates are varying from country to country. This is because of the climate variability, which is known to affect the outbreaks of many infectious diseases. Thus different infection fatality rate might be taking place by the impacts of environmental factors on the genomic content of SARS-CoV-2, changing the functionality or virulence at the particular condition. Consequently more investigations would be required for better investigation of SARS-CoV-2 genomic architecture and the hereditary varieties as indicated by different ecological states of various nations. The study aims to unravel the reason for the different infection and variable death rates amongst the aftected countries in the world.

Advances in next-generation sequencing (NGS) has resulted in an unprecedented explosion of genomic sequence data. The genome-wide association studies (GWAS) and comprehensive identification of genetic variants in the form of single nucleotide polymorphisms (SNPs) across the genomes of individuals is a definite prerequisite for translational genomic medicine ^9-11^. Thus this study focuses on the genomic sequence divergence and genetic variability within the available whole genome sequences of SARS-CoV-2 isolated from affected countries with different infction mortality rates.

## 2. Material and Methods

### 2.1. Sequence Mining

There were total 92 whole-genome sequences of SARS-CoV-2 virus from different preceding countries throughout the world, have been deposited in the genome database (https://www.ncbi.nlm.nih.gov/genome/) of National Centre for Biotechnology Information (NCBI) (https://www.ncbi.nlm.nih.gov/) web portal. Among these available genome sequences, this study retrieved a total of 11 genome sequences of first cases of SARS-CoV-2 isolated from 11 different counties, including the reference genome isolated from Wuhan, China, in FASTA format. Along with the genome sequence of SARS-CoV-2, the genome sequence of Bat Coronavirus has also collected from NCBI for further phylogeny study.

### 2.2. Multiple Sequence Alignment and Phylogenetic Analysis

All the collected genome sequences of SARS-CoV-2 were subjected to multiple sequence alignment through the Molecular Evolutionary Genetics Analysis 7 (MEGA 7)^12^ (https://www.megasoftware.net/) package by using the MUSCLE algorithm. Among the aligned sequences, the highly mismatched sequences or the large gap regions have removed. The good quality based aligned sequences have considered for generating the phylogenetic tree by using the maximum parsimony method with 1000 bootstrap replicates.

### 2.3. Study of Nucleotide Diversity and Haplotype

The study of genomics diversity has been implemented on the aligned sequences to analyze the difference between the genome sequences of SARS-CoV-2 by examining the variable sites, Nucleotide, and haplotype diversity in retrieved genome sequences through DNA sequence polymorphism software DNAsp 6 (http://www.ub.edu/dnasp/) ^13^. The resulted haplotype diversity has represented through a population genetic software, i.e., population analysis with reticulate trees (PopART 1.7) (http://popart.otago.ac.nz/index.shtml) ^14^.

### 2.4. Isolation of Mutation Sites in Coding Regions

The resulted variable sites have been analysed, and identified the respective nucleotides in 11 genome sequences from GenBank database (https://www.ncbi.nlm.nih.gov/genbank/) of NCBI. The respective coded genes in the variable sites have also been analyzed.

### 2.5. Retrieval of Worldwide Reports of COVID19

The populations by 2020 of analysed countries have been collected from world meter (https://www.worldometers.info/world-population/population-by-country/). Also, the total confirmed cases and number of deaths of COVID-19 as per 6th June 20202 had collected from WHO. These retrieved data have been used to calculate the infection fatality rate of COVID-19 in selected countries. Further, the co-relation impact of mutational effects on death rates has been analysed.

## 3. Results

### 3.1. Sequence Analysis

The collected whole-genome sequences of SARS-CoV-2 isolated from 11 different countries, including Brazil, Nepal, USA, Australia, South Korea, Taiwan, Sweden, India, Italy, Vietnam, and China (Table 1) have aligned through MEGA. Including 11 sequences of SARS-CoV-2, the genome sequence of Bat coronavirus has also been aligned to represent sequence divergence by constructing a phylogenetic tree followed by maximum parsimony method with 1000 bootstrap replicates. The established phylogenetic tree contains 2 clades, Clade I and II. In Clade I, the genome sequences of the USA, Taiwan, Nepal, Vietnam, China, and India have been grouped, but India has diversified with individual taxa from the other countries. However, in Clade II, Sweden, Italy, Australia, South Korea, and Brazil have been grouped. Meanwhile, the genome sequence of Bat coronavirus clearly showed its divergence from another 11 genome sequence of SARS-CoV-2, as it has not included in any of the resulting clades of the phylogenetic tree (Fig. 1).

**Table 1:**
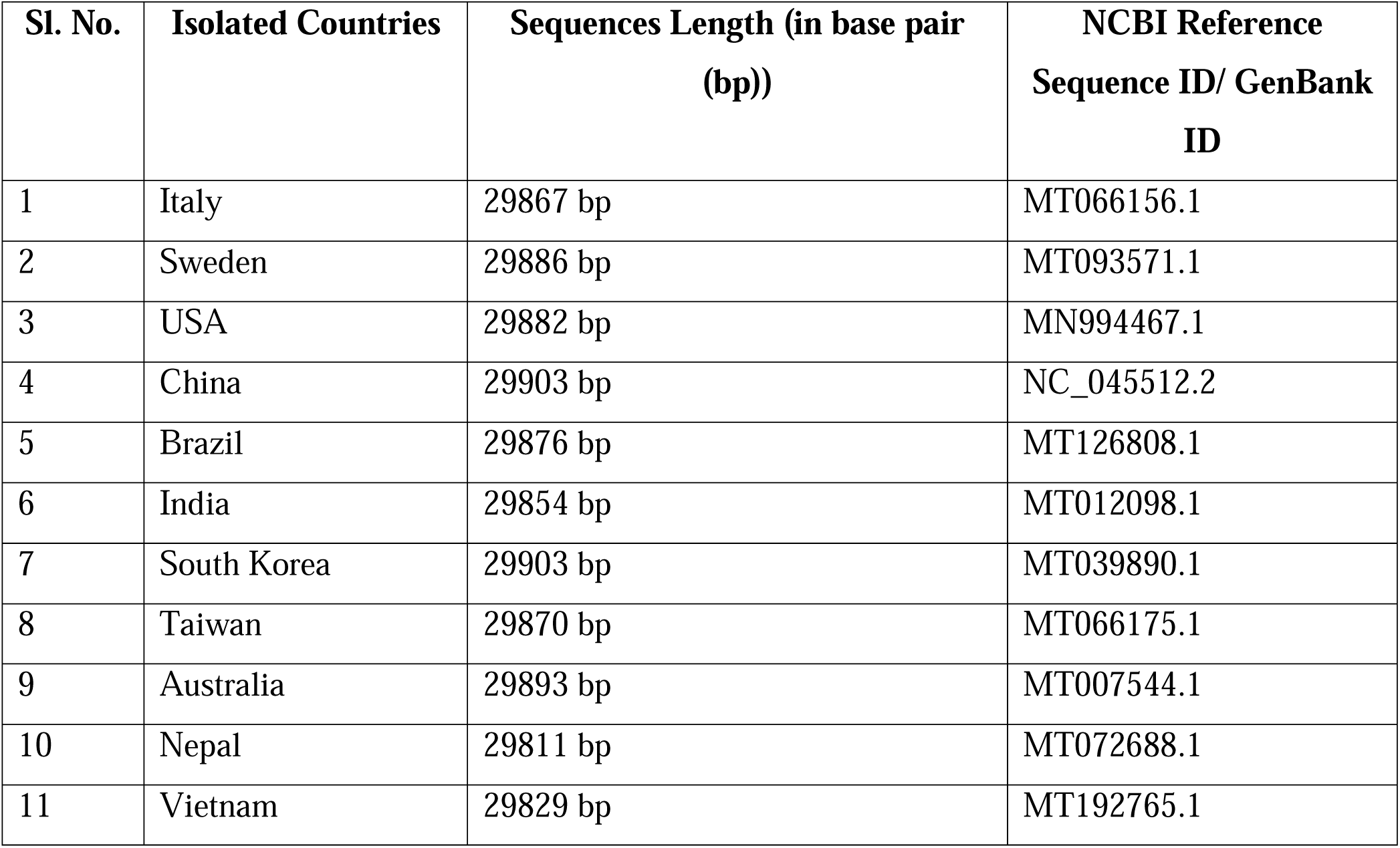
Collected Genome Sequences of 1^st^ cases of SARS-CoV-2, isolated from 11 countries

| Sl. No. | Isolated Countries | Sequences Length (in base pair (bp)) | NCBI Reference Sequence ID/ GenBank ID |
| --- | --- | --- | --- |
| 1 | Italy | 29867 bp | MT066156.1 |
| 2 | Sweden | 29886 bp | MT093571.1 |
| 3 | USA | 29882 bp | MN994467.1 |
| 4 | China | 29903 bp | NC_045512.2 |
| 5 | Brazil | 29876 bp | MT126808.1 |
| 6 | India | 29854 bp | MT012098.1 |
| 7 | South Korea | 29903 bp | MT039890.1 |
| 8 | Taiwan | 29870 bp | MT066175.1 |
| 9 | Australia | 29893 bp | MT007544.1 |
| 10 | Nepal | 29811 bp | MT072688.1 |
| 11 | Vietnam | 29829 bp | MT192765.1 |

**Fig. 1:**
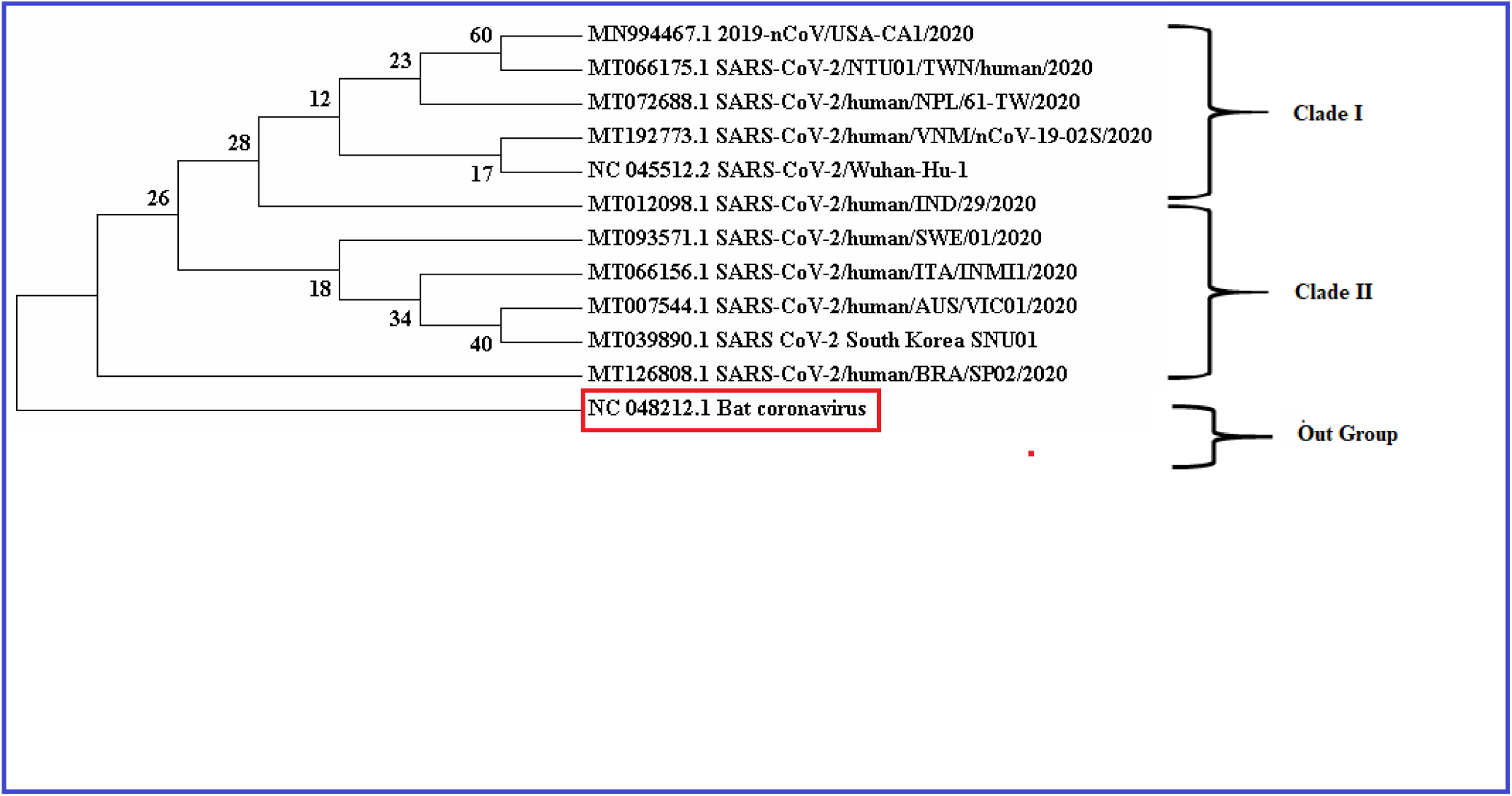
Phylogenetic tree represents the diversified whole genome sequences of SARS-CoV-2, isolated from 11 different countries with bootstrap frequencies for each taxon

### 3.2. Diversity Study

In the multiple aligned genome sequences, the nucleotide diversities and/or DNA polymorphism have analyzed through DNAsp. There were a total of 29990 nucleotide sides, with 0.00023 nucleotide diversity, which has resulted. The polymorphic site analysis has predicted 29762 monomorphic (invariable) sites, 34 polymorphic (variable) sites, or mutations. Meanwhile, the haplotype diversity has resulted in total 11 haplotypes with 1.000 haplotype diversity within 11 whole-genome sequences of SARS-CoV-2 (Table 2). The haplotype network represents the number of genetic mutations between the sequences (Fig. 2).

**Table 2:** Resulted DNA Polymorphisms and Polymorphic Sites in SARS-CoV-2 Genome Sequences

| DNA Polymorphism |  | Polymorphic Data |  |
| --- | --- | --- | --- |
| No. of Polymorphic Sites (S) | 34 | Polymorphic Sites:<br>Invariable (Monomorphic) Sites | 29762 |
|  |  | Variable (Polymorphic) Sites | 34 |
| Total No. of Mutations (Eta) | 34 | No. of Singleton Variable Sites | 30 |
| No. of Haplotypes (h) | 11 | Singleton Variable Sites | 28, 1548, 2269, 2277, 2717, 2971, 6031, 6695, 9274, 10232, 12115, 13225, 13226, 14657, 14805, 15597, 17373, 17376, 19065, 20936, 22224, 22303, 22785, 23952, 25775, 26354, |
|  |  |  | 26729, 28077, 28792 |
| Haplotype Diversity (hd) | 1.000 | No. of Parsimony Variable Sites | 4 |
| No. of Nucleotide Sites | 29990 | Parsimony Variable Sites | 8782, 24034, 26144, 28144 |
| Nucleotide Diversity (Pi) | 0.00023 |  |  |

**Fig. 2:**
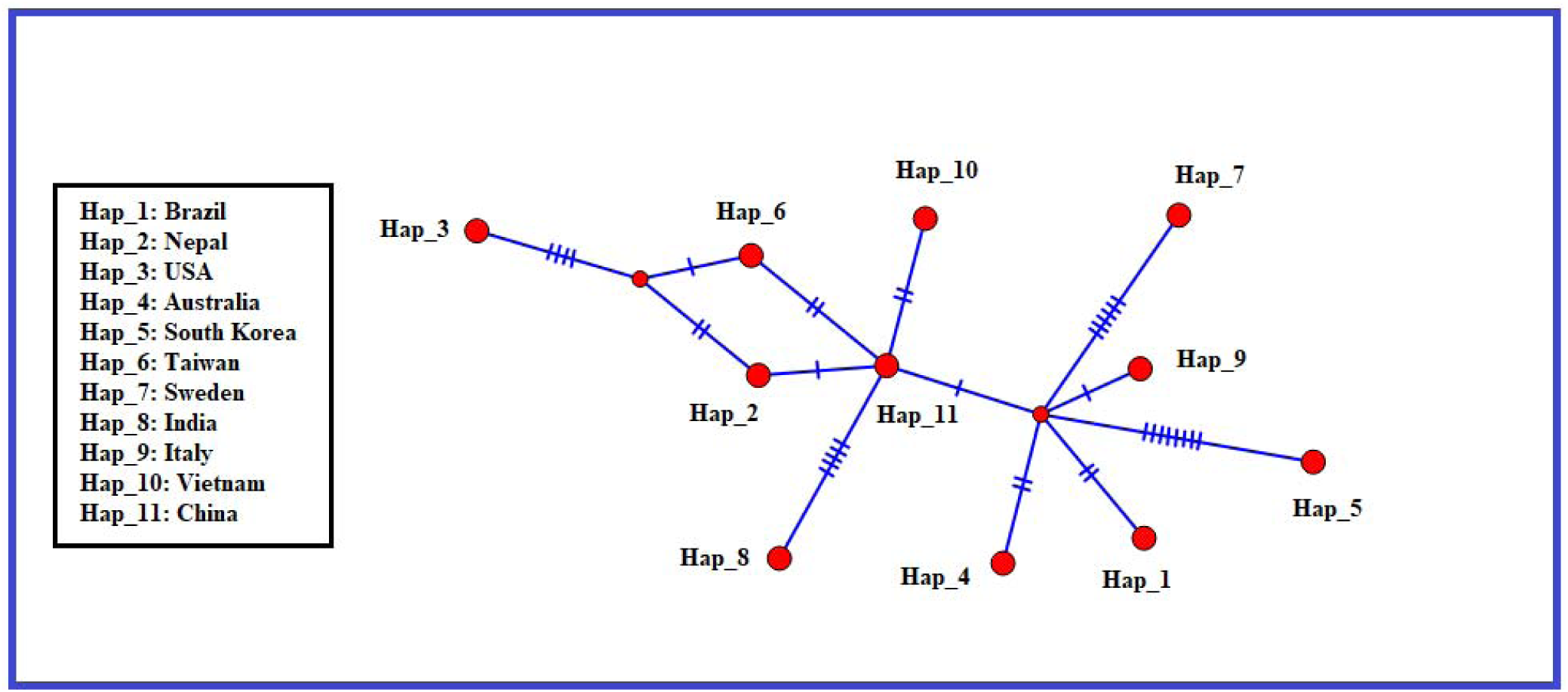
Presented integer neighbor joining method based haplotype network with number of mutations with in the genome sequences of SARS-CoV-

### 3.3. Mutational analysis

On analysis, 34 variable regions or polymorphic sites were identified, out of which 30 were singletons i.e. 28, 1548, 2269, 2277, 2717, 2971, 6031, 6695, 9274, 10232, 12115, 13225, 13226, 14657, 14805, 15597, 17373, 17376, 19065, 20936, 22224, 22303, 22785, 23952, 25775, 26354, 26729, 28077, 28792 and 4 were parsimony sites i.e. 8782, 24034, 26144, 28144. (Table 3). All the variable sites have been analyzed in GenBank to isolate the respective nucleotides for all 11 genome sequences to identify the single nucleotide mutations/polymorphisms (SNPs) in these particular variable sites (Table 3). According to the analyzed SNPs, there were no mutations found in singleton sites 22785 and 23952. In contrast, the most sequence variations with three nucleotides found in singleton variable regions, including 2971, 6031, and 10232 in 11 countries. However, some mutations contained the complementary base pair nucleotides.

**Table 3:** Analyzed Single Nucleotide Mutations in the Resulted Variable Regions

| Variable Positions | Brazil | Nepal | USA | Australia | Korea | Taiwan | Sweden | India | Italy | Vietnam | Ch |
| --- | --- | --- | --- | --- | --- | --- | --- | --- | --- | --- | --- |
| **Singleton** |  |  |  |  |  |  |  |  |  |  |  |
| 28 | C | T | C | C | C | C | C | T | T | T | T |
| 1548 | G | G | A | G | G | G | G | T | G | G | G |
| 2269 | A | G | A | A | A | A | A | G | T | A | A |
| 2277 | T | A | T | T | T | T | T | T | T | T | T |
| 2717 | G | G | G | G | G | G | A | A | G | G | G |
| 2971 | G | A | G | G | T | G | G | T | G | G | G |
| 6031 | C | T | C | C | T | C | C | A | C | C | C |
| 6695 | C | G | C | C | C | C | C | A | C | C | C |
| 9274 | A | T | A | A | A | A | G | A | A | A | A |
| 10232 | C | A | C | C | C | C | C | A | C | T | C |
| 12115 | C | A | C | C | T | C | C | G | C | C | C |
| 13225 | C | G | C | C | C | C | G | T | C | C | C |
| 13226 | T | T | T | T | T | T | C | C | T | T | T |
| 14657 | C | A | C | C | C | C | C | C | C | C | C |
| 14805 | T | A | C | C | C | C | C | C | C | C | C |
| 15597 | T | A | T | T | C | T | T | C | T | T | T |
| 17373 | C | G | C | C | C | C | C | T | C | C | C |
| 17376 | A | T | A | A | A | A | G | A | A | A | A |
| 19065 | T | A | T | C | T | T | T | G | T | T | T |
| 20936 | C | C | C | C | T | C | C | T | C | C | C |
| 22224 | C | T | C | C | G | C | C | G | C | C | C |
| 22303 | T | T | T | G | T | T | T | G | T | T | T |
| 22785 | G | G | G | G | G | G | G | G | G | G | G |
| 23952 | T | T | T | T | T | T | T | T | T | T | T |
| 25775 | G | G | G | G | T | G | G | C | G | G | G |
| 26354 | T | A | T | T | A | T | T | C | T | T | T |
| 26729 | T | T | C | T | T | T | T | T | T | T | T |
| 28077 | G | T | C | G | G | G | G | C | G | G | G |
| 28792 | A | C | T | A | A | A | A | G | A | A | A |
| **Parsimony** |  |  |  |  |  |  |  |  |  |  |  |
| 8782 | C | T | T | C | C | T | C | A | C | C | C |
| 24034 | C | A | T | C | C | C | C | G | C | C | C |
| 26144 | T | T | G | T | T | G | T | T | T | G | G |
| 28144 | T | A | C | T | T | C | T | T | T | T | T |

Furthermore, these identified variable regions of SARS-CoV-2 genome sequences have defined as protein-coding regions. This study has analyzed only the coding regions of Spike (S) protein and Nucleocapsid (N) protein in the genome sequence of SARS-CoV-2 of 11 countries submitted at NCBI. It has identified that the coding regions of S (21563-25384) and N protein (28274-29533) of SARS-CoV-2 are similar in nine countries from four different continents. The coding regions of S protein and N protein in genome sequences obtained from India [S (21550-25368) and N (28258-29517)] and Nepal [S (21548-25369) and N (28274-29533)] are different from rest sequences isolated from other countries. However, the resulted variable sites within the coding regions of S protein are 22224, 22303, 22785, 23952 and the variable site within N protein of SARS-CoV-2 is 28792.

### 3.4. SNPs in Protein Coding Sites

#### a. S protein coding region variability

The found SNPs in the S protein coding region 22224 are variable between India, Korea, and Nepal i.e. at this position “G” is present in the genome of SARS-CoV-2 isolated from India and South Korea; “T” is present in Nepal; whereas rest of the isolated genomes from other countries contain “C” at 22224 position. Likewise, the isolated genome sequence of SARS-CoV-2 from India and Australia at position 22303 contains “G,” which varies from different countries. There are no mutations found in 22785 and 23952 positions. (Table 3)

#### b. N protein coding region variability

In the coding variable position 28792 of N protein India, Nepal and USA showed variation than other countries as follows; Indian genome sequence contains “G”, in case of Nepal the present Nucleotide is “C” and in case of USA the Nucleotide is “T” and in other countries the genomic sequence contains “A” at the respective position, which codes N protein of SARS-CoV-2. (Table 3)

### 3.5. Analysis of Infection Fatality Rate

The infection fatality rates have calculated from the retrieved number of confirmed cases of COVID-19 and deaths related to COVID-19 as per 6^th^ July, 2020 of the pandemic in 11 different countries (Fig. 3). According to the resulted fatality rates, Italy (0.144) has the highest death rate, followed by Sweden (0.075). In Asian countries, China has (0.054) the highest case fatality rate compared to India (0.028), South Korea (0.021), Taiwan (0.015), and Nepal (0.002). Interestingly country like Vietnam (0) has a zero fatality rate of COVID-19 till 6th July 2020 (Fig. 4). Two significant countries from both the North and South American continent, i.e., USA (0.0456) and Brazil (0.040) have comparable infection fatality rate. (Table 4). The fatality rate for Australia was 0.012. When comparing deaths and the number of cases based on population for the USA, approximately 0.85% of the population has been infected, while in India, only 0.05% of the population has the disease (Table 4) (Fig. 3).

**Table 4:** Collected Cases, Deaths and Calculated Case Fatality Rates of COVID19 in Different Countries as per 6^th^ July, 2020

| Sl. No. | Countries | Population (2020) | No. of Cases (As per 6<sup>th</sup> July 2020) | No. of Deaths (As per 6<sup>th</sup> July 2020) | Calculated Case Fatality Rates (Total Deaths/ Total Cases) | %age of Infection on the basis of total Population (Total Cases/Total Population*100) |
| --- | --- | --- | --- | --- | --- | --- |
| 1 | Italy | 60,461,826 | 241,611 | 34,861 | 0.144 | 0.399% |
| 2 | Sweden | 10,099,265 | 71,419 | 5,420 | 0.075 | 0.007% |
| 3 | USA | 331,039,958 | 2,833,552 | 129,408 | 0.045 | 0.855% |
| 4 | China | 1,439,430,327 | 85,320 | 4,648 | 0.054 | 0.005% |
| 5 | Brazil | 212,588,506 | 1,577,004 | 64,265 | 0.040 | 0.741% |
| 6 | India | 1,380,159,707 | 697,413 | 19,693 | 0.028 | 0.050% |
| 7 | South Korea | 51,269,185 | 13,181 | 285 | 0.021 | 0.025% |
| 8 | Taiwan | 23,816,775 | 449 | 7 | 0.015 | 0.001% |
| 9 | Australia | 25,499,884 | 8,449 | 104 | 0.012 | 0.033% |
| 10 | Nepal | 29,136,808 | 15,784 | 34 | 0.002 | 0.054% |
| 11 | Vietnam | 97,355,481 | 369 | 0 | 0 | 0.000% |

**Fig. 3:**
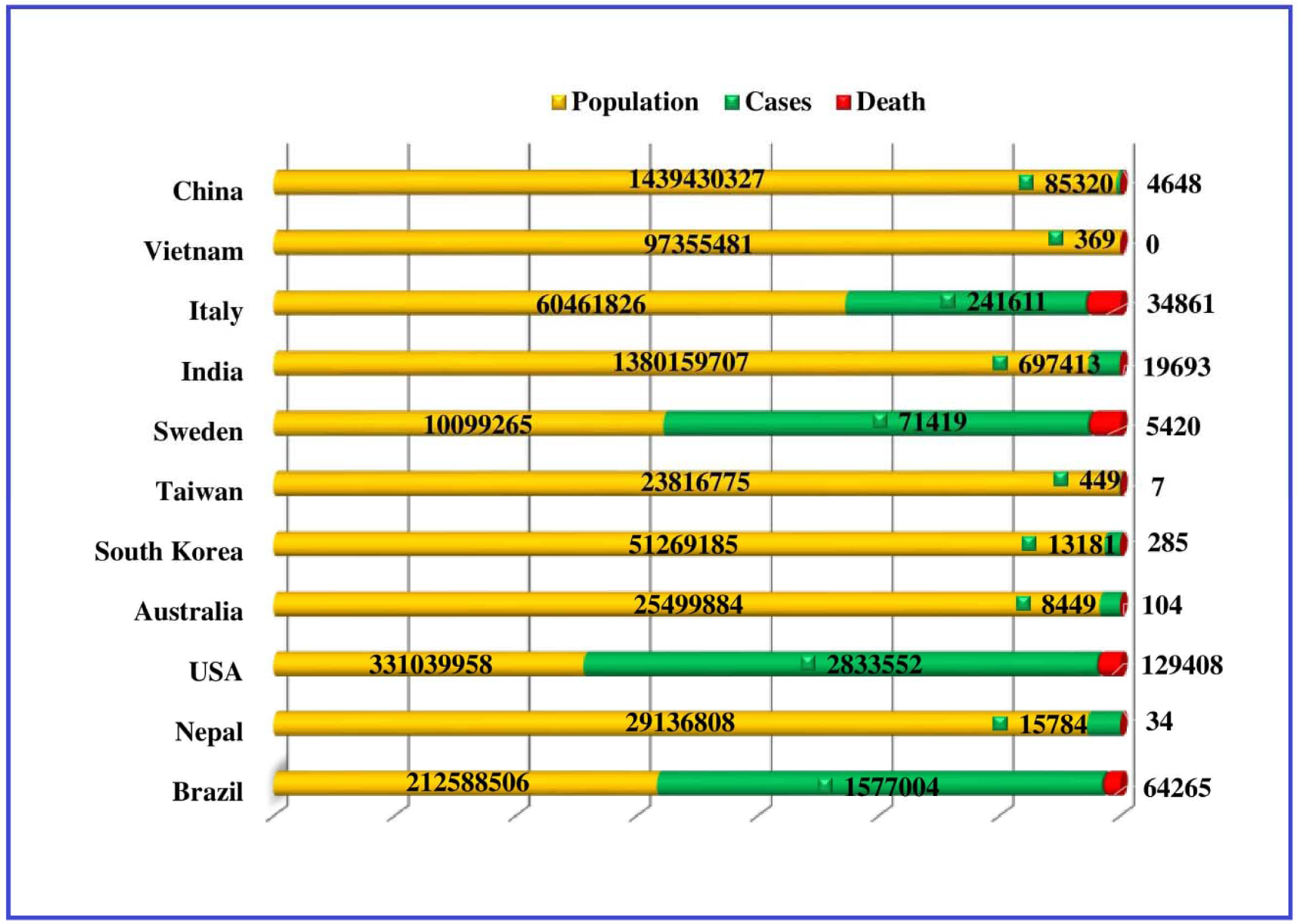
Bar chart schematically shows the total population of 11 different countries, with number of cases, and deaths of COVID-19 as per 6^th^ July, 2020 collected from WHO

**Fig. 4:**
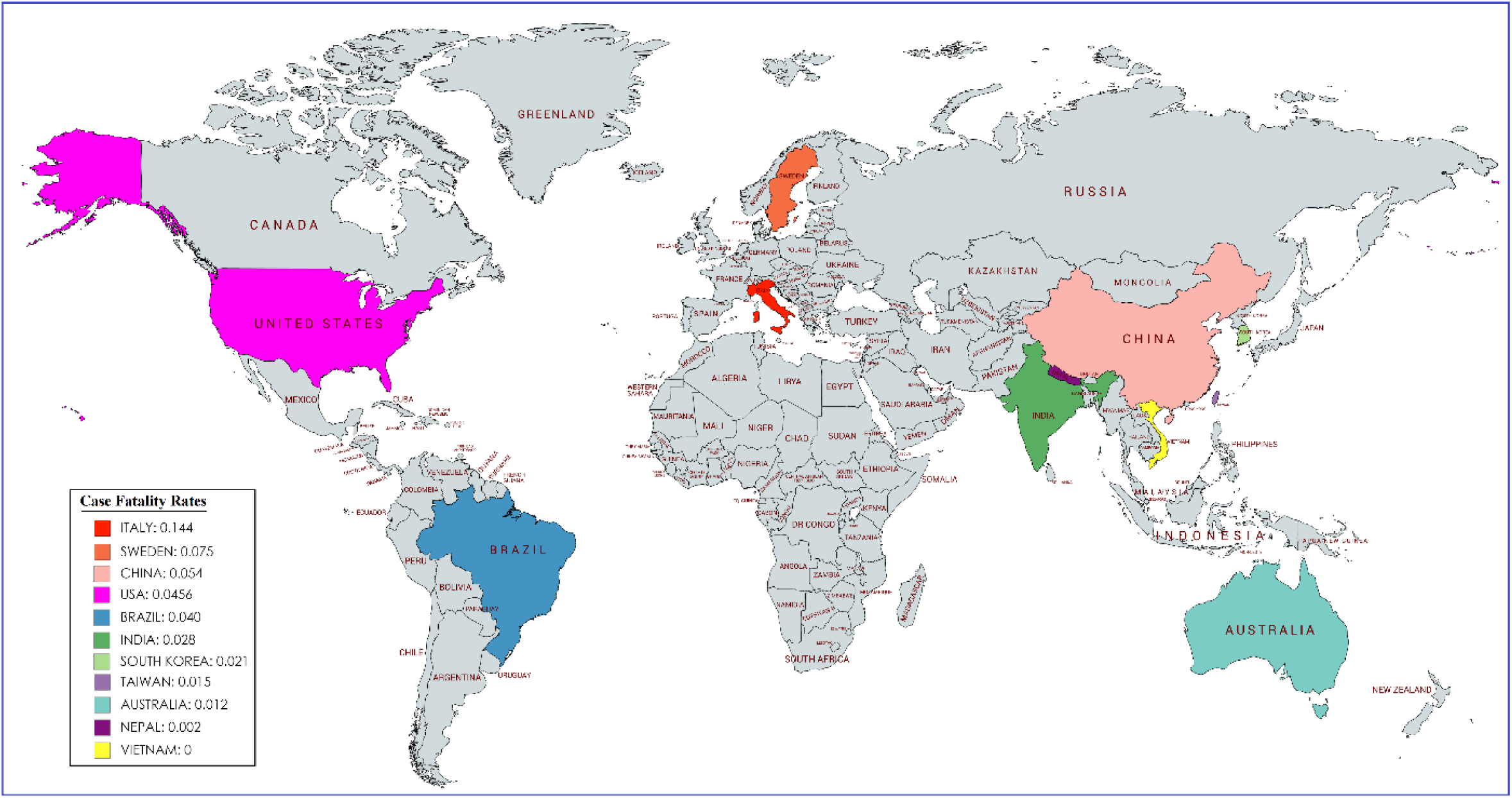
Calculated case fatality rates for 11 different countries represented in world map with specific color tags

## 4. Discussion

The fast transmission of novel SARS-CoV-2, among individuals in the various areas of the world, is apparently keeping pandemic unrestricted. The spread of this emerging virus is mainly by droplets, which is affecting the respiratory tract mucosa and attaches with ACE receptor present of the surface epithelium of the alveoli of lungs. Because of its exceptional genomic development, it has a higher transmission rate than SARS and MERs, and it is amazingly testing to characterize the job of hereditary changeability as well as polymorphisms in the genomic content separated from various nations.

The constructed phylogenetic tree represents the closely related and divergence in the genomic sequence of SARS-CoV-2, isolated from 11 countries. In the two major clades, I and II Nepal, India, and Brazil have diverged from other sequences that represent the genetic diversity in the SARS-CoV-2 genome sequences isolated from these regions. These outcomes have more clarified by the genetic diversity study, including nucleotide assorted variety and polymorphism investigation, which are the key boundaries to recognize the genomic fluctuation or arrangement contrasts as SNPs ^15-16^ by comparing the nucleotide sequences. SNPs are the most common type of genetic variations and have become an essential genetic marker for human mapping disease, genetics, and evolutionary studies ^15^. The resulted 11 haplotypes in the aligned genome sequences showed the number of mutations and the genetic difference between the genomic alleles of SARS-Cov-2, although in the analyzed 34 polymorphic sites altered nucleotides have found which satisfied the differentiation in isolated genome sequences of SARS-CoV-2. Meanwhile, according to S protein-coding regions, India, Korea, Nepal showed the altered Nucleotide than others and in case of N protein-coding regions SNPs found in the SARS-CoV-2 genome sequences isolated from India and Australia. Again it has been identified that India and Nepal have different protein coding regions in comparison with other countries, which has also been represented in phylogenetic tree. So from the above perceptions it has been clearly verified that India has the most diversified SARS-CoV-2 as it is having altered nucleotides at the viral protein coding positions and also having different genomic sequence than others. After India, Nepal might be harbouring the 2nd most diverse species of SARS-CoV-2.

According to the previous research data, GWAS has been used vigorously in the field of genetic variation studies and SNP analysis. The nearness of SARS-CoV-2 genomic information empowers scientists for more investigation and examination on it, for example, the hereditary variation examination and SNP seclusion, nucleotide varieties at the genomic positions have been analyzed between the COVID-19 cases and control samples of Italian and Spanish cohorts ^17^; between the symptomatic and asymptomatic COVID-19 severity patients^18^; some genomic mutational sites have been also identified in Europe and North America^19^; also the complete genome sequences of SARS-CoV-2 have been studied to understand the genomic structures and the most common mutations within^20^. Apart from these the haplotype network analysis has also been done on the genomic contents of Japanese people^21^. Before the isolation of SARS-CoV-2, the nucleotide sequences of isolated SARS-CoV had compared and analyzed to identify the positions of SNPs based on the genomic polymorphism ^22^. Also, the MERS-CoV genomes were aligned and phylogenetically analyzed to identify the SNP variants ^23^.

The current investigation explores the potential varieties among the entire genome sequences of SARS-CoV-2, isolated from 11 different countries. The observed results in the present study has identically urged some novel hereditary data including the nucleotide assorted varieties, SNPs, mutations in particular protein coding districts of SARS-CoV-2 species. The mutations might effected in the functional variability of viral genes or the virulence factors of SARS-CoV-2, which reflects on the case fatality rates and mortality rates of respective countries. These findings might be used in the development of a vaccine or drug candidates against COVID-19 based on the different genomic architecture of SARS-CoV-2 in a greater extent. They might also be able to predict the course of the disease or disease pattern in a particular country or region of the world.

## Conclusion

The whole world currently tangled in developing a novel vaccine against SARS-CoV-2. Meanwhile, the above investigation is not just restricted to the genomic varieties, but also gives the mutational consequences for protein-coding regions, which additionally be used for the improvement of proper immunization candidates according to the genomic variability to conquer the current world pandemic circumstances. This study might also be used as a prediction tool for the disease behavior and expected infection fatality rate in a region or a country of the world. Based on this study, the various countries may prepare or brace themselves by gauging on the upcoming health care tsunami they are about to face.

